# Recombination and repetitive genomic landscapes are decoupled in a close relative of *Caenorhabditis elegans*

**DOI:** 10.64898/2026.05.30.728836

**Authors:** Kimberly A. Moser, Courtney King, Gavin C. Woodruff

## Abstract

Genomes exhibit chromosomal heterogeneity. Distributions of genes, repetitive elements, and polymorphisms are not uniform along chromosomes in multiple species. One explanation for these patterns is recombination rate variation. As recombination interacts with selection to shape the evolutionary fates of alleles, recombination rate variation could promote differences in the chromosomal distribution of genomic features. Thus, clarifying the relationship between recombination rates and genomic organization is a major goal of genetics. In the nematode *Caenorhabditis elegans*, recombination rate correlates with multiple genomic features that are non-uniformly distributed along chromosomes; recombination rates and repeats are higher on chromosome ends compared to chromosome centers. Its closest known relative, *C. inopinata*, harbors a radically altered genome with nearly uniform chromosomal distributions of repetitive elements. Is this dramatic change in genomic organization connected to the evolution of recombination rates? Here, we describe a genetic map of *C. inopinata* constructed via whole-genome sequencing of 180 individual F_2_ recombinants. This reveals four chromosomes have a conserved recombination rate domain structure whereas two other chromosomes harbor divergent, more uniform recombination rate distributions. Comparisons of these intrachromosomal recombination rates with genomic features reveal little covariation between recombination rate, diversity, gene density, and repeat content in *C. inopinata* (in stark contrast to most *Caenorhabditis* species). This suggests that the evolution of recombination may not be entirely responsible for the atypical uniform distribution of repetitive elements across *C. inopinata* chromosomes. Taken together, these observations reveal that recombination rates can be decoupled from the genomic organization of repetitive elements.

## Introduction

Genomes are organized. That is, the various features constituting genomes are not randomly or uniformly distributed across chromosomes. These features are diverse and include repetitive elements (Maxwell 2020; Kejnovsky et al. 2025); coding genes (Keller et al. 2000; Hou et al. 2012; Sehgal et al. 2014); polymorphisms (Burri et al. 2015; Nelson and Cresko 2018; Harringmeyer and Hoekstra 2022); patterns of population differentiation (Poelstra et al. 2014; Nelson and Cresko 2018); non-coding genes (Olena and Patton 2010; Kawaoka et al. 2013); heterochromatic and euchromatic regions (Janssen et al. 2018); and high chromatin-chromatin contact regions (van Steensel and Belmont 2017; Falk et al. 2019); among others. Large-scale chromosomal features, such as centromeres, drive the evolution of some such patterns of genomic organization (Schueler and Sullivan 2006; Barra and Fachinetti 2018; Muller et al. 2019). Likewise, the existence of genetic sex determination (and the sex chromosomes associated with such systems) carries outsized influence on the evolution of genomic organization (Graves 2008; Zhu et al. 2025). Moreover, in some systems, dramatic inversions carrying alleles under strong selection appear to drive the organization of multiple genomic features (Hoffmann and Rieseberg 2008; Harringmeyer and Hoekstra 2022; Westram et al. 2022). And, the existence of ancient, highly conserved gene cassettes such as the *Hox* cluster suggest that some measure of genomic organization is maintained by strong selection in segmented animals ((Ruddle et al. 1994); although some animals, like octopi, have retained *Hox* genes on separate chromosomes while losing their clustered organization (Seo et al. 2004)). It is then clear that there are many factors driving genomic organization.

Despite this, centromeres cannot explain the chromosomal distributions of all genomic features, as genomes can harbor heterogeneity in non-centromeric regions. Moreover, genomic structure persists on holocentric chromosomes (Senaratne et al. 2022). Along the same lines, autosomes can have intrachromosomal genomic organization, so sex chromosomes cannot explain such organization entirely. Many species do not harbor population-specific insertions, and most genes do not seem to cluster together on chromosomes by molecular or physiological function (John and Miklos 1988). And, the magnitude and direction of genomic organization varies tremendously by species and the genomic feature under consideration (Lynch 2007). Thus, the myriad causes of genomic organization remain unresolved.

Recombination is thought to be a major contributor to genomic organization. As a shuffler of haplotypes, recombination can both unite beneficial alleles and free such alleles from detrimental neighboring polymorphisms (Charlesworth and Jensen 2021). Conversely, the absence of recombination is thought to impede adaptation by keeping novel beneficial alleles in linkage disequilibrium with deleterious variants (Muller 1964; Hartfield and Keightley 2012). Thus, variation in recombination rates along chromosomes could impact the efficacy of selection *across* chromosomes, potentially affording a satisfying explanation of genomic organization (Nachman 2002; Lynch 2007). Indeed, recombination rates are frequently non-uniform across chromosomes (McVean et al. 2004; Stapley et al. 2017). And, recombination rates correlate with many genomic features. Recombination rate covaries with intraspecific polymorphisms in many systems, suggestive of widespread linked selection across eukaryotic systems (Charlesworth and Campos 2014; Ellegren and Galtier 2016; Kern and Hahn 2018; Jensen et al. 2019). And, one of the major lessons of the high-throughput sequencing era appears to be the pervasiveness of linked selection (Charlesworth and Jensen 2021), making recombination a central subject of contemporary evolutionary genomics (Henderson and Bomblies 2021; Johnston 2024).

Repetitive elements have likewise been of great interest within molecular evolution for decades (McClintock 1950; Britten and Kohne 1968). Dominating eukaryotic genomes (Wells and Feschotte 2020), the pervasiveness of simple tandem repeats, as well as more complex transposable elements, holds no one straightforward and intuitive explanation. Transposable elements in particular are a subject of fascination as selfish genetic elements, revealing eukaryotic genomes as arenas of genomic conflict and antagonistic coevolution churning at the molecular scale of biological organization (Werren 2011; Ågren and Clark 2018). In general, transposable elements are thought to be deleterious as they can insert into coding sequences (disrupting protein function), and transposable elements are then expected to accumulate in regions of low recombination where selection is less effective (Lippman et al. 2004; Haddrill et al. 2007; Betancourt et al. 2009). Empirical patterns generally hold up to these theoretical expectations, with transposable element density positively correlated with recombination rate in most systems (Fig. 1a; (Kent et al. 2017)). However, notable differences have been observed when considering transposable element types. Transposable elements harbor variable replication strategies, with type II transposons having a “cut-and-paste” transposition mode, whereas type I retrotransposons reproduce via a “copy-and-paste” mechanism (Wicker et al. 2007). For instance, whereas LINE/SINE elements in rice (Tian et al. 2009) and Type II elements in wheat (Daron et al. 2014) reveal a positive correlation with recombination rate, these elements reveal a negative correlation with recombination rate in fruit flies (Rizzon et al. 2002) and rats (Jensen-Seaman et al. 2004). Recombination, selection, and mode of transposable element replication may then interact to shape patterns of repetitive genomic organization.

**Figure 1.**
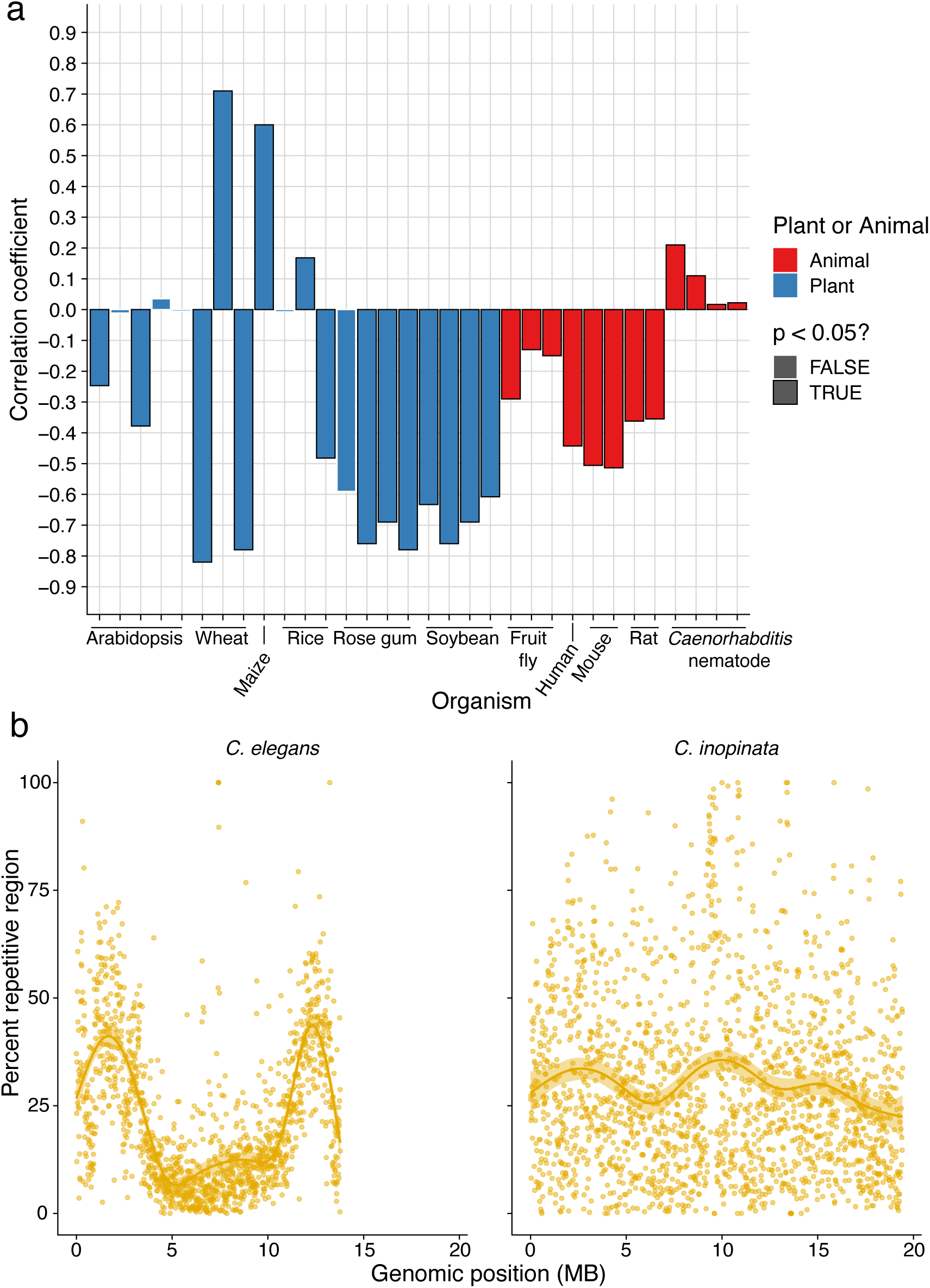
Repetitive element densities are correlated with recombination rates. (a) Previous data reveal recombination rate and repeat content are positively correlated in *Caenorhabditis* nematodes, despite being negatively correlated in many organisms (Data from Kent et al. 2017 Figure 1; Woodruff and Teterina 2020; and this study). (b) Whereas *C. elegans* harbors a typical arm-enriched repetitive genomic landscape, *C. inopinata* maintains a more uniform distribution of repetitive elements along chromosomes (Data from Woodruff and Teterina 2020).

Yet, there are some organismal systems that contradict the theoretical and conventional empirical empirical observations. *Caenorhabditis* nematodes harbor an atypical repetitive genomic landscape wherein repetitive elements accumulate in *high* recombining regions (THE C. ELEGANS SEQUENCING CONSORTIUM 1998; Stein et al. 2003; Fierst et al. 2015; Teterina et al. 2020; Woodruff and Teterina 2020; Noble et al. 2021; Stevens et al. 2022; Sun et al. 2022; Teterina et al. 2023). However, *Caenorhabditis* nematodes, like other systems, reveal notable correlations among genomic features with respect to their chromosomal organization. Genes are typically enriched in low-recombining chromosome centers (THE C. ELEGANS SEQUENCING CONSORTIUM 1998; Stein et al. 2003; Fierst et al. 2015; Kanzaki et al. 2018; Teterina et al. 2020; Noble et al. 2021; Stevens et al. 2022; Sun et al. 2022; Teterina et al. 2023). Polymorphisms are enriched in high-recombining chromosome arms (Andersen et al. 2012; Thomas et al. 2015; Noble et al. 2021; Teterina et al. 2023; Woodruff et al. 2024). Heterochromatin is likewise partitioned to less transcriptionally active chromosome arms (Liu et al. 2011). These patterns of gene density and chromatin type may influence repeat density across chromosomes (Charlesworth and Langley 1989; Lippman et al. 2004). In addition, the existence of holocentric centromeres, wherein chromosomes synapse across their entire lengths, may also contribute to the atypical relationships between repeat density and recombination in nematodes (Senaratne et al. 2022). Regardless, it remains unclear exactly why these nematodes appear to harbor a repetitive genomic landscape that contradicts the expectations of evolutionary theory (Barrón et al. 2014) and diverges from broadly observed patterns in other organisms (Fig 1a; (Kent et al. 2017)).

*Caenorhabditis inopinata* harbors a repetitive genomic landscape that is highly divergent from its close relatives (Fig. 1b; (Woodruff and Teterina 2020)). Instead of harboring TE-enriched chromosome arms, TEs are uniformly distributed on *C. inopinata* chromosomes (Fig. 1b; (Woodruff and Teterina 2020)). We previously found that this pattern was largely driven by a handful of TE superfamilies with diverse replication mechanisms (including both Type I and Type II elements; (Woodruff and Teterina 2020)). *C. inopinata* retains conserved, gene-enriched chromosome centers, although the magnitude of this effect is diminished in this species (Woodruff and Teterina 2020). A similar pattern holds for intraspecific genetic diversity — in *C. inopinata*, diversity is enriched on chromosome arms, but to a smaller degree than observed in *C. elegans* (Woodruff et al. 2024). Regardless, previous evolutionary simulations we performed revealed that (unsurprisingly) uniform distributions of recombination rates across chromosomes promote uniform distributions of TEs (Woodruff and Teterina 2020). Recombination rates evolve (Henderson and Bomblies 2021), and some large-effect mutations in *C. elegans* can homogenize chromosome-wide recombination rates (Zetka and Rose 1995; Parée et al. 2024). Thus, it is possible that the evolution of recombination rates in *C. inopinata* drives the evolution of its genomic organization.

However, the distribution of recombination rates across chromosomes has not been measured in *C. inopinata*. Here, we describe the generation of a genetic map to infer chromosome-wide estimates of recombination rates in *C. inopinata*. We compare these observations with previously-published genetic maps (Rockman and Kruglyak 2009; Noble et al. 2021; Stevens et al. 2022; Teterina et al. 2023; Parée et al. 2024) to understand the evolution of genomic organization in *Caenorhabditis* nematodes. We find that the extent of chromosomal recombination rate heterogeneity has been diminished in *C. inopinata*, with two autosomes revealing nearly uniform recombination rate distributions. Thus, the recombination rate landscape has been decoupled from the repetitive genomic landscape in this species, suggestive that the evolution of recombination rates is not the only driver of genome-wide TE evolution in this species.

## Materials and methods

### Strains and culture conditions

The recombinant *C. inopinata* individuals used to create the genetic map were derived from the parental strains NKZ35 and WOU9. NKZ35 is a ten-generation inbred line derived from the first wild isolate of this species from Ishigaki Island, Okinawa (NKZ2/NK74SC); this line was used to generate the *C. inopinata* reference genome (Kanzaki et al. 2018). WOU9, an inbred line, was generated following ten generations of sibling pair inbreeding; this line was derived from a *C. inopinata* wild isolate from Taiwan (PX723) (Hammerschmith et al. 2022). Animals were maintained on Nematode Growth Media (Brenner 1974) made with 3.2% agar. Plates were seeded with *E. coli* OP50 as a food source. Animals were grown at 25°C as *C. inopinata* exhibits better reproduction at higher temperatures compared to *C. elegans* (Kanzaki et al. 2018; Woodruff et al. 2019).

### Generation of recombinants

The parental lines *C. inopinata* NKZ35 and WOU9 were mated in both cross directions. F_1_ siblings were crossed to generate 180 total F_2_ recombinants (90 F_2_ recombinants from each reciprocal cross were sequenced). Four maternal NKZ35 F_2_ recombinants were ultimately excluded from downstream analyses due to having a low fraction of reads aligning to the *C. inopinata* reference genome (<70%). F_2_ adults were washed three times in Phosphate Buffered Saline (PBS). Individual animals were then transferred to individual tubes filled 10 μl of TE with 5% Proteinase K. Single worms were incubated for one and a half hours at 60°C for lysis with a subsequent incubation at 95°C incubation for 15 minutes for enzyme inactivation. In addition to recombinant F_2_ animals, individuals from the two parental lines, as well as F1 individuals, were also prepared for whole-genome sequencing. Single-worm DNA preparations were then used for whole-genome sequencing.

### Library preparation and sequencing

Linear amplification of single-worm DNA (5 μl of the crude preparation above) was performed with the Illustra GenomiPhi V3 amplification kit (GE Lifesciences). DNA was purified with the Zymo Genomic DNA Clean and Concentrator kit. DNA then was readied for whole-genome sequencing via a miniaturized Nextera library preparation approach using an Echo liquid handler; 150 bp paired-end reads were generated with the Illumina Novaseq 6000 S4 platform (University of Oregon GC3F Genomics Core Facility).

### Read processing, alignment, and genotyping

Read quality was assessed with *FastQC* and *MultiQC* (Ewels et al. 2016) using the default parameters. Reads were aligned to the *C. inopinata* reference assembly (retrieved from WormBase ParaSite (Howe et al. 2017) version PRJDB5687.WBPS18) with *bwa mem* (version 0.7.13) using default parameters (Li 2013). Alignment files were sorted and indexed with *samtools* (version 1.11) *sort* and *samtools index* using default parameters (Danecek et al. 2021).

Parental SNPs appropriate for genetic mapping were defined as those (1) homozygous and shared within each parental line and (2) never shared between parental lines. Parental alignment files were merged with *samtools merge* (with default parameters) to generate single alignment files for each parental line (NKZ35 and WOU9). Multiply-mapping reads were excluded via SAM FLAG tags (using the command “*grep-v-e’XA:Z:’-e’SA:Z:’*”). Genotypes were called with *bcftools* (version 1.11) *mpileup* (with options “-Ou”) and *bcftools call* (with options “-c-Ob”) (Danecek et al. 2021). Sites with <10x coverage were excluded (with *bcftools view* options “-i’DP>=10’”). Parental BCF files were merged with *bcftools merge* and biallelic SNPs were extracted with *bcftools view* (options “-m2-M2-v snps”). The Bash functions *awk* and *sed* were then used to identify sites with alternative homozygous states in NKZ35 and WOU9. This yielded 49,821 sites with potential biallelic SNPs differentiating the two parental strains. As the *C. inopinata* genome is highly repetitive (Kanzaki et al. 2018; Woodruff and Teterina 2020), an attempt was made to exclude such regions. The hard-masked version of the *C. inopinata* genome (version PRJDB5687.WBPS18) was used to generate a BED file of presumptive repetitive regions (using the python script *generate_masked_ranges.py*; https://www.danielecook.com/generate-masked-ranges-bedfile-fasta/). The function *bedtools complement* was then used to generate a list of non-repetitive sites (Quinlan and Hall 2010); *bedtools intersect* was subsequently used to define the sites harboring fixed differences among the parental lines in non-repetitive regions (yielding 41,442 sites). To aid in genetic map inference, *in silico* F_1_ samples were generated by randomly sampling an equal number of reads from NKZ25 and WOU9 parental.bam files (for males, reads from the X were sampled from only one parent).

For mapping populations, PCR duplicates were inferred and removed with *picard MarkDuplicates* (GATK version 4.5.0.0; with option “REMOVE_DUPLICATES=true”); read groups were added with *picard AddOrReplaceReadGroups*; and alignment files were indexed with *picard BuildBamIndex* with default parameters (https://broadinstitute.github.io/picard/). Base calls were calibrated with *picard BaseRecalibrator* (with the above parental SNPs masked with the option “--known-sites’) and *picard ApplyBQSR*. Multiply-mapping reads were subsequently excluded as described above. Sites defining the parental SNPs characterized above were extracted from recombinant samples with *samtools view* (options “-F 4-bS-q 15-L parental_snp_sites.bed”). Alignment files were sorted and indexed with *samtools* (version 1.11) *sort* and *samtools index* using default parameters (Danecek et al. 2021). Coverage across relevant sites was inferred with *bedtools coverage*, and one low-coverage sample (“F2A-29”) was excluded from downstream analyses.

### Inference of C. inopinata genetic map

Genotypes and recombination maps were inferred with Lep-MAP3 (Rastas 2017). Genotypes for each chromosome were inferred with *samtools mpileup* (options “-q 10-Q 10-s”) and the Lep-MAP3 function *Pileup2Likelihoods* with default parameters. For each chromosome, the Lep-MAP3 function *ParentCall2* (with options “posteriorFile=- removeNonInformative=1”) was used to call parental genotypes. Genotypes for each chromosome were then passed to the Lep-MAP3 function *Filtering2* (with options “dataTolerance=0.001”) to filter markers exhibiting potential segregation distortion. Filtered markers and genotypes were then passed to the Lep-MAP3 function *OrderMarkers2* to infer genetic maps for each chromosome (autosome options: “data=- improveOrder=0 sexAveraged=1 grandparentPhase=1”; X chromosome option: “data=-improveOrder=0 recombination1=0 grandparentPhase=1”). For visualizing genotype frequencies (Figure 2b), genotypes were extracted with the Lep-MAP3 script *map2genotypes.awk* (“awk-vfullData=1-f *map2genotypes.awk*”).

**Figure 2.**
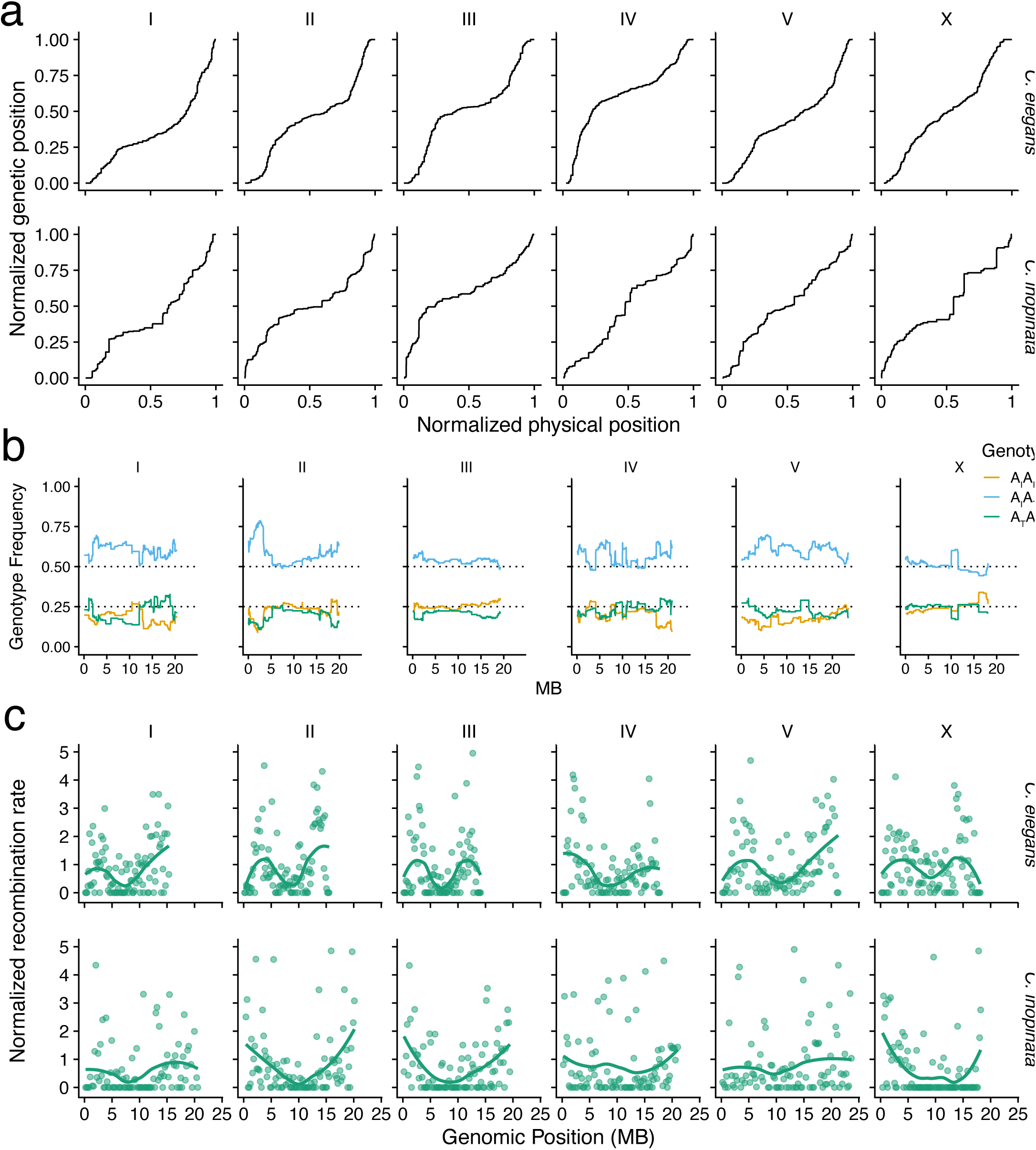
Chromosomal patterns of recombination are partially conserved between *C. elegans* and *C. inopinata*. (a) Marey maps for *C. elegans* (Data from Rockman and Kruglyak 2009) and *C. inopinata* (this study). (b) Genotype frequencies among *C. inopinata* F2 recombinants (A_I_ = Ishigaki NKZ35 allele; A_T_ = Taiwan WOU9 allele). (c) Patterns of normalized recombination rates across physical genomic spans binned by chromosome. Each point represents a rank ordered one-hundredth non-overlapping chromosomal bin.

### Comparative genomics

To enable comparisons among *Caenorhabditis* species, previously generated genetic maps for *C. elegans* (Rockman and Kruglyak 2009; Parée et al. 2024), *C. briggsae* (Stevens et al. 2022), *C. remanei* (Teterina et al. 2023), and *C. tropicalis* (Noble et al. 2021) were retrieved. Gene count and repeat density data for *C. elegans*, *C. inopinata*, *C. remanei*, and *C. briggsae* were retrieved from a previous study on repetitive genomic landscapes (Woodruff and Teterina 2020). For *C. tropicalis*, gene annotations and repetitive regions were retrieved from a separate study (Noble et al. 2021). Previously-generated intraspecific genetic diversity data were likewise retrieved from previous studies on *C. elegans* (Noble et al. 2021), *C. inopinata* (Woodruff et al. 2024), *C. briggsae* (Noble et al. 2021), *C. remanei* (Teterina et al. 2023), and *C. tropicalis* (Noble et al. 2021). For comparing genomic features across species, recombination rates, gene counts, genetic diversity metrics, and repeat densities were normalized by chromosome length. This was done by partitioning each species-specific chromosome into one hundred equally-sized bins and then determining the mean values (recombination rate, gene count, etc.) per bin. Associations among recombination rates and genomic features (Fig. 4-5) were performed with ordinary least-squares regression with corrections for multiple comparisons with the Benjamini-Hochberg method. For defining chromosome arms and centers, arms were defined as bins ranked <25 and> 75; centers were defined as bins ranking from 25-75. These are the values used in Fig. 3-7. To correlate recombination rates with repeat types, the repeat taxonomy and density data from (Woodruff and Teterina 2020) was retrieved, normalized as above, and joined with the recombination rate data. Ordinary least squares regression was performed for repeat types, *p*-values were corrected for multiple comparisons via the Benjamini-Hochberg method, and significant associations were plotted (Fig. 8). The following R packages were used for analyses and visualizations: *cowplot* (Wilke 2020), *dplyr* (Wickham, François, et al. 2023), *ggforce* (Pedersen 2022a), *ggplot2* (Wickham 2016), *ggstats* (Larmarange 2026), *lemon* (Edwards 2020), *patchwork* (Pedersen 2022b), *rstatix* (Kassambara 2023), and *tidyr* (Wickham, Vaughan, et al. 2023).

**Figure 3.**
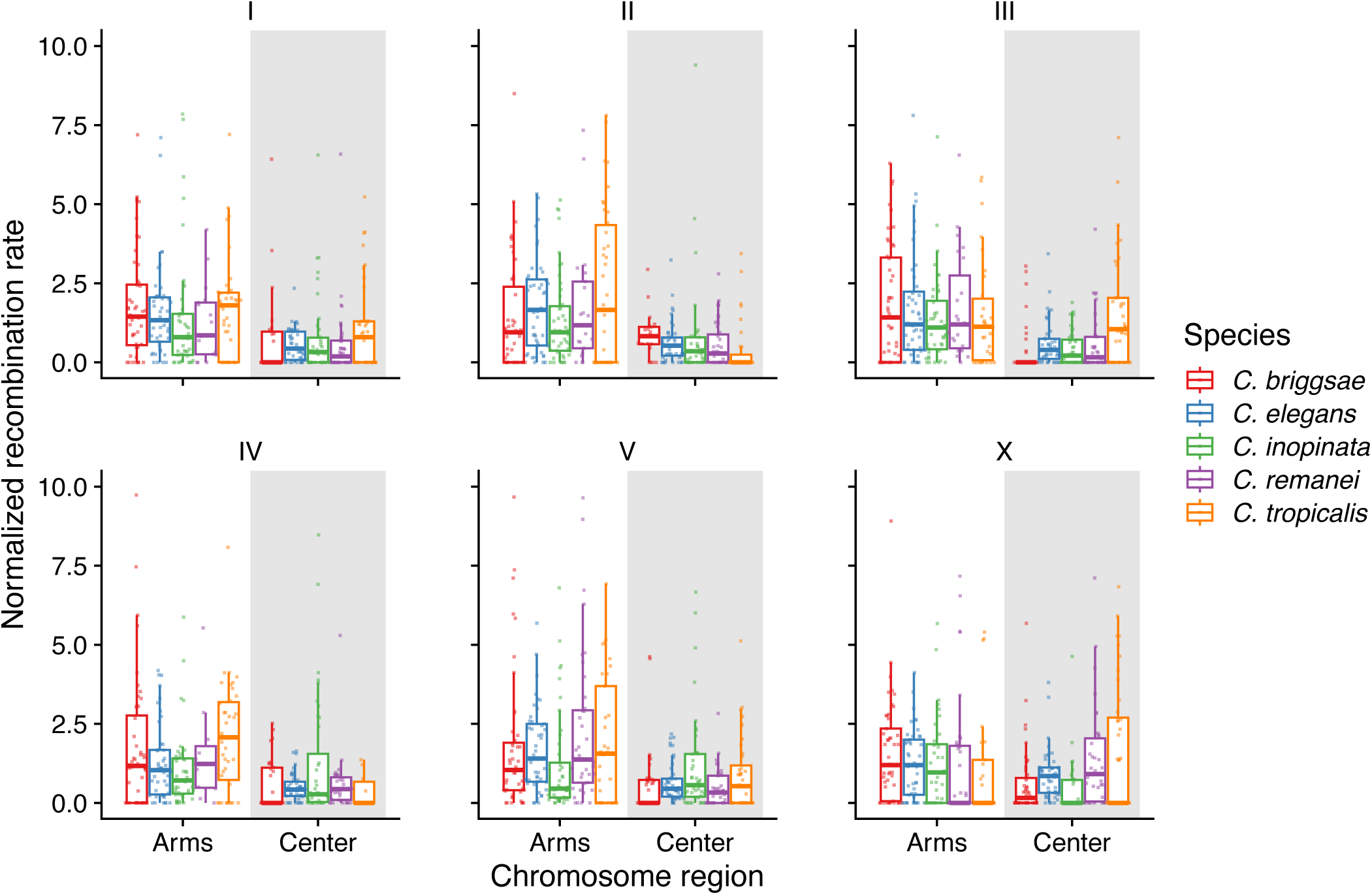
Recombination rates are elevated in chromosome arms compared to chromosome centers across *Caenorhabditis* species. Recombination rate data (Rockman and Kruglyak 2009; Noble et al. 2021; Stevens et al. 2022; Teterina et al. 2023; this study) were retrieved and normalized by chromosome position. Center halves are defined as chromosome centers; outer halves are defined as chromosome arms. Each point represents a rank ordered one-hundredth non-overlapping chromosomal bin.

**Figure 4.**
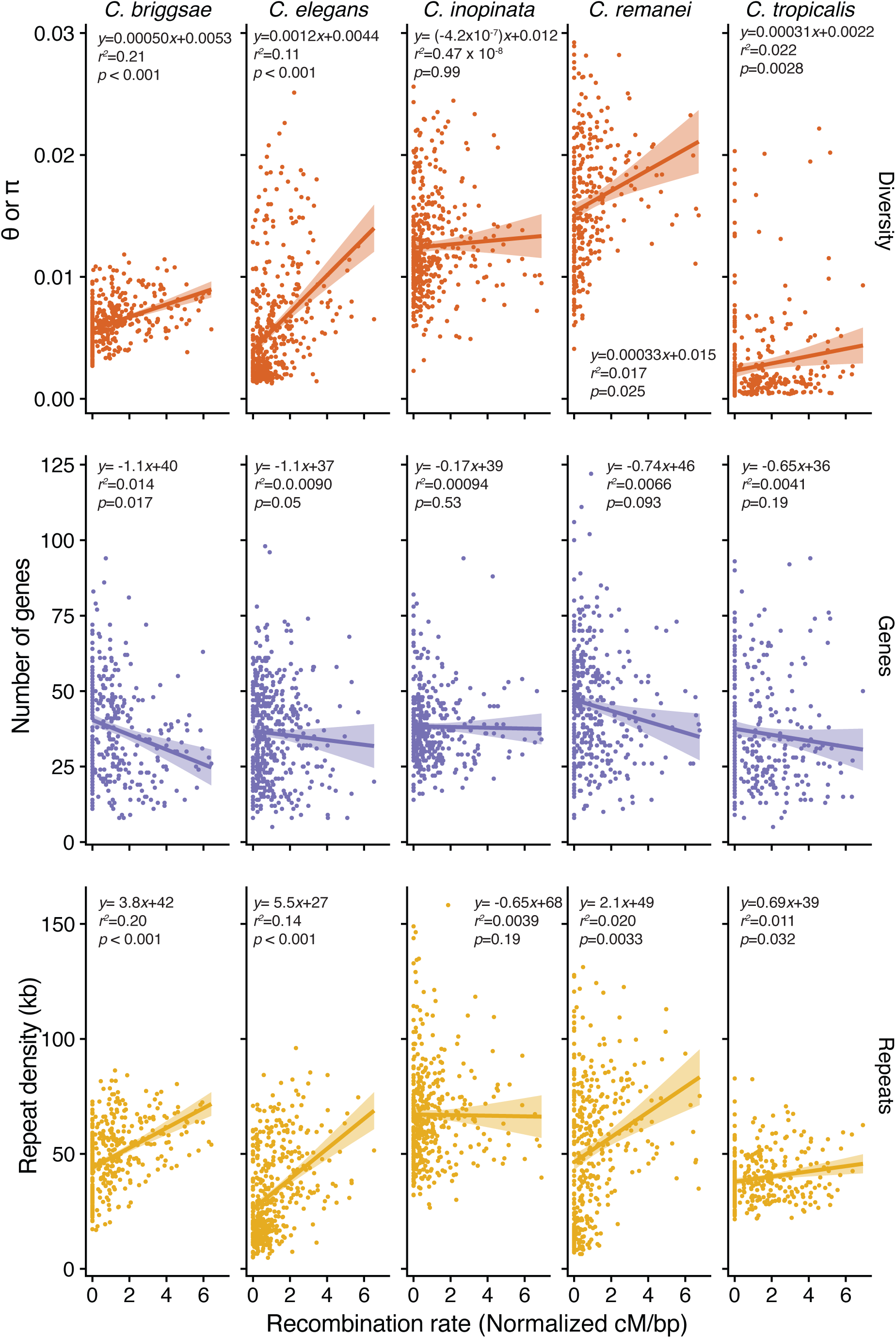
Recombination rate, repeat density, gene content, and genetic diversity are largely decoupled in *C. inopinata*. Recombination rate data (Rockman and Kruglyak 2009; Noble et al. 2021; Stevens et al. 2022; Teterina et al. 2023; this study) were retrieved and normalized by chromosome position. Diversity data represents estimates of Watterson’s θ in *C. briggsae* (Noble et al. 2021), *C. tropicalis* (Noble et al. 2021), *C. elegans* (Noble et al. 2021) and *C. remanei* (Teterina et al. 2023) or nucleotide diversity in *C. inopinata* (Woodruff et al. 2024). Gene and repeat density data were retrieved from previous studies (Woodruff and Teterina 2020; Noble et al. 2021). Ribbons represent 95% confidence intervals. Each point represents a rank ordered one-hundredth non-overlapping chromosomal bin.

**Figure 5.**
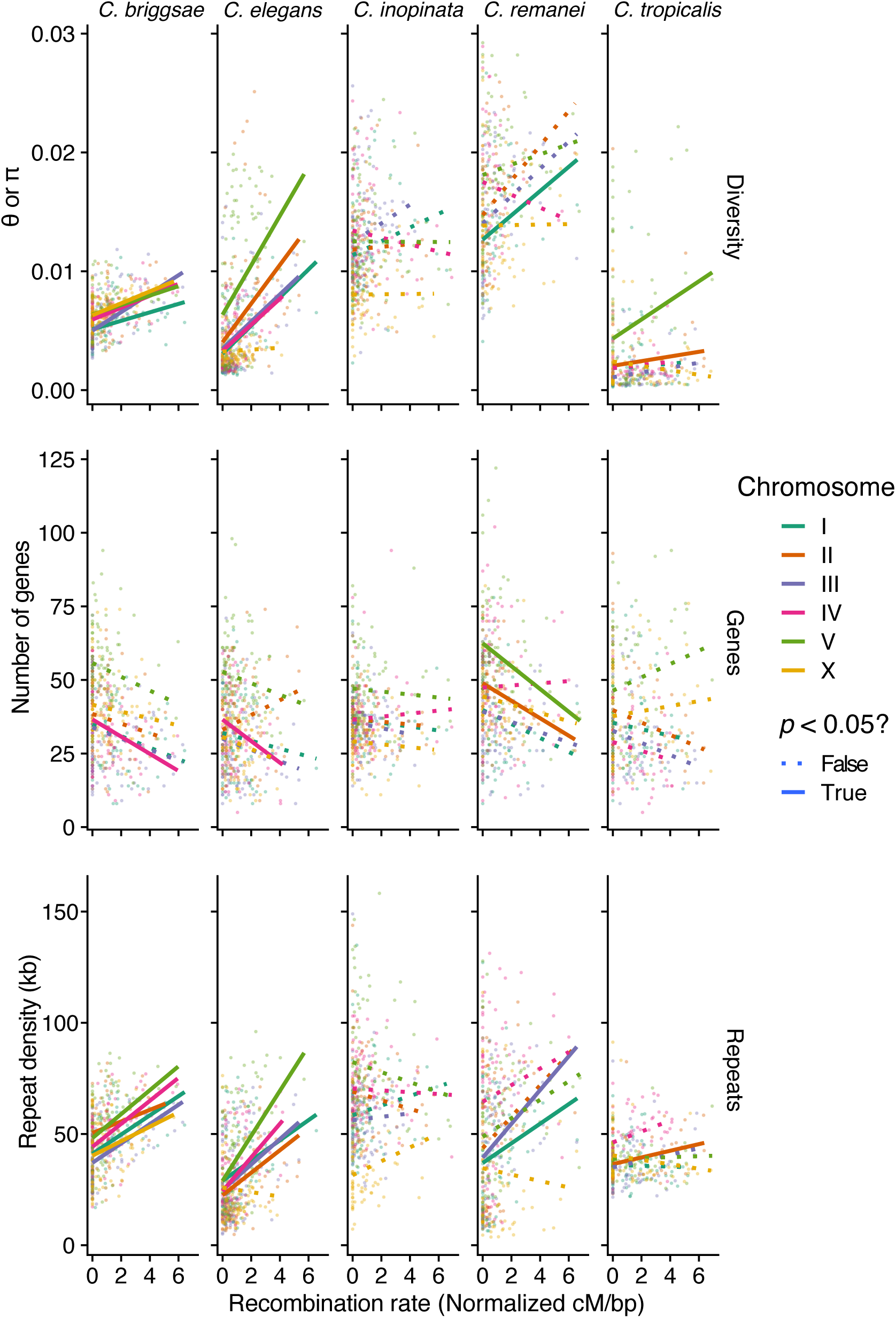
Recombination rate and genomic feature correlations vary by chromosome. Data plotted in Figure 4 are colored by chromosome. Solid lines represent linear models with FDR-adjusted *p*-values < 0.05.

**Figure 6.**
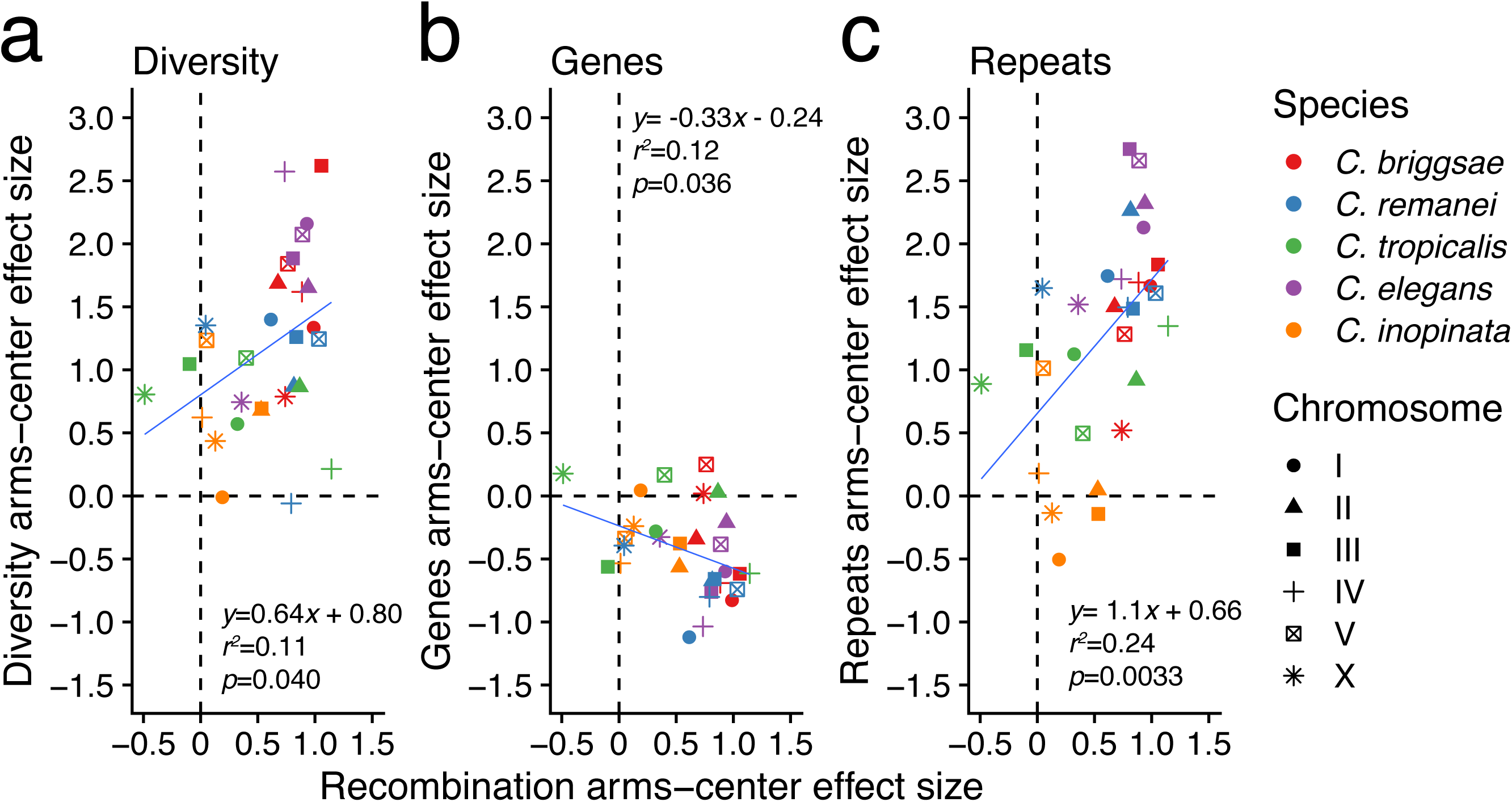
Chromosomal differentiation in recombination is correlated with differentiation in diversity gene density, repeat density, and in *Caenorhabditis*. Plotted are Cohen’s *d* effect sizes among chromosome arms and centers for each feature per chromosome. Values of 0 reveal no differences among chromosome arms and centers. Positive values reveal features enriched on chromosome arms. Negative values reveal features enriched on chromosome centers. Diversity (a), Gene density (b), and repeat density (c) effect sizes are plotted on the y-axes.

**Figure 7.**
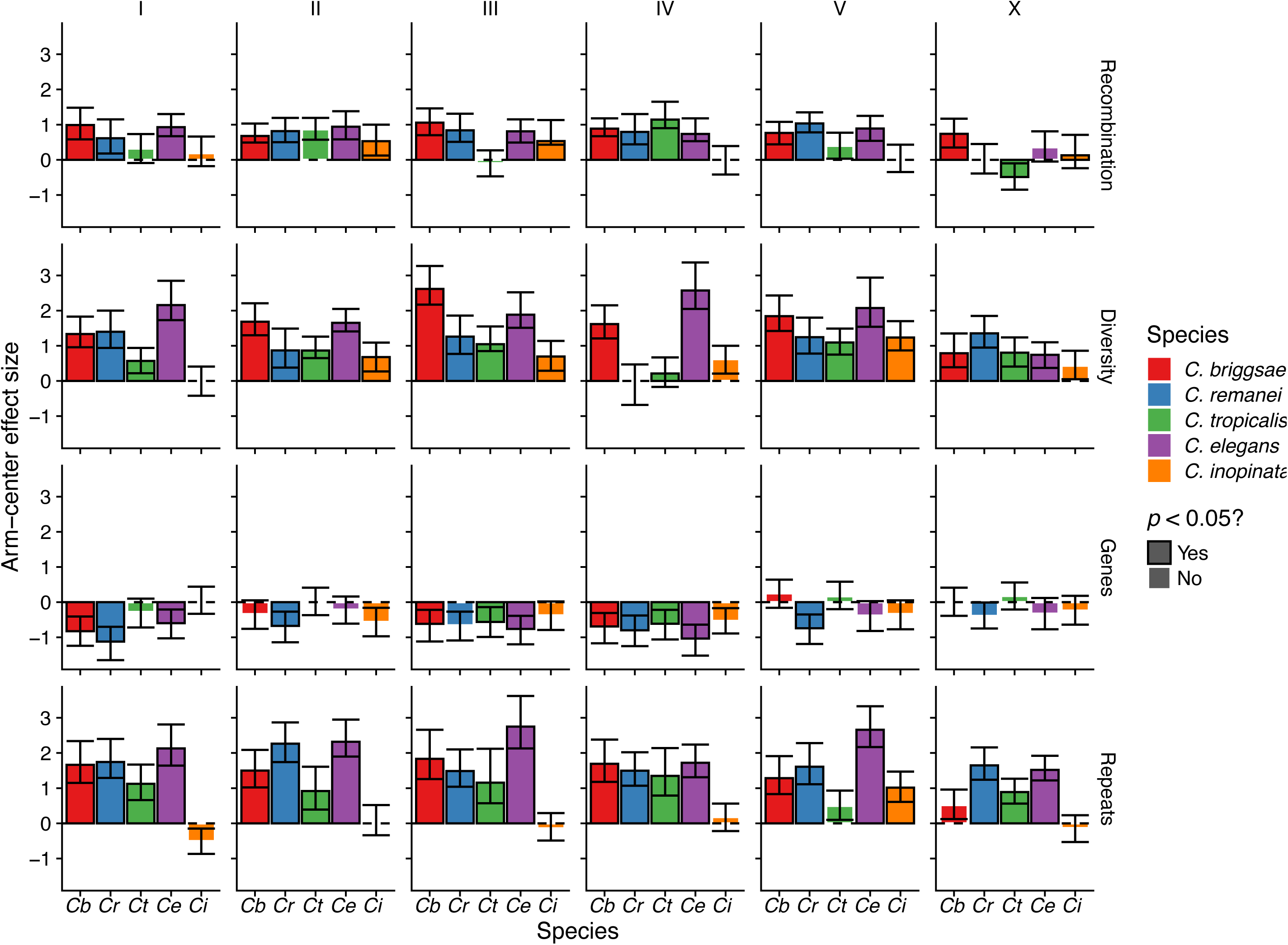
Arm-center effect sizes for all genomic features and chromosomes across five *Caenorhabditis* species. Plotted are Cohen’s *d* effect sizes among chromosome arms and centers for each feature per chromosome. Values of 0 reveal no differences among chromosome arms and centers. Positive values reveal features enriched on chromosome arms. Negative values reveal features enriched on chromosome centers.

**Figure 8.**
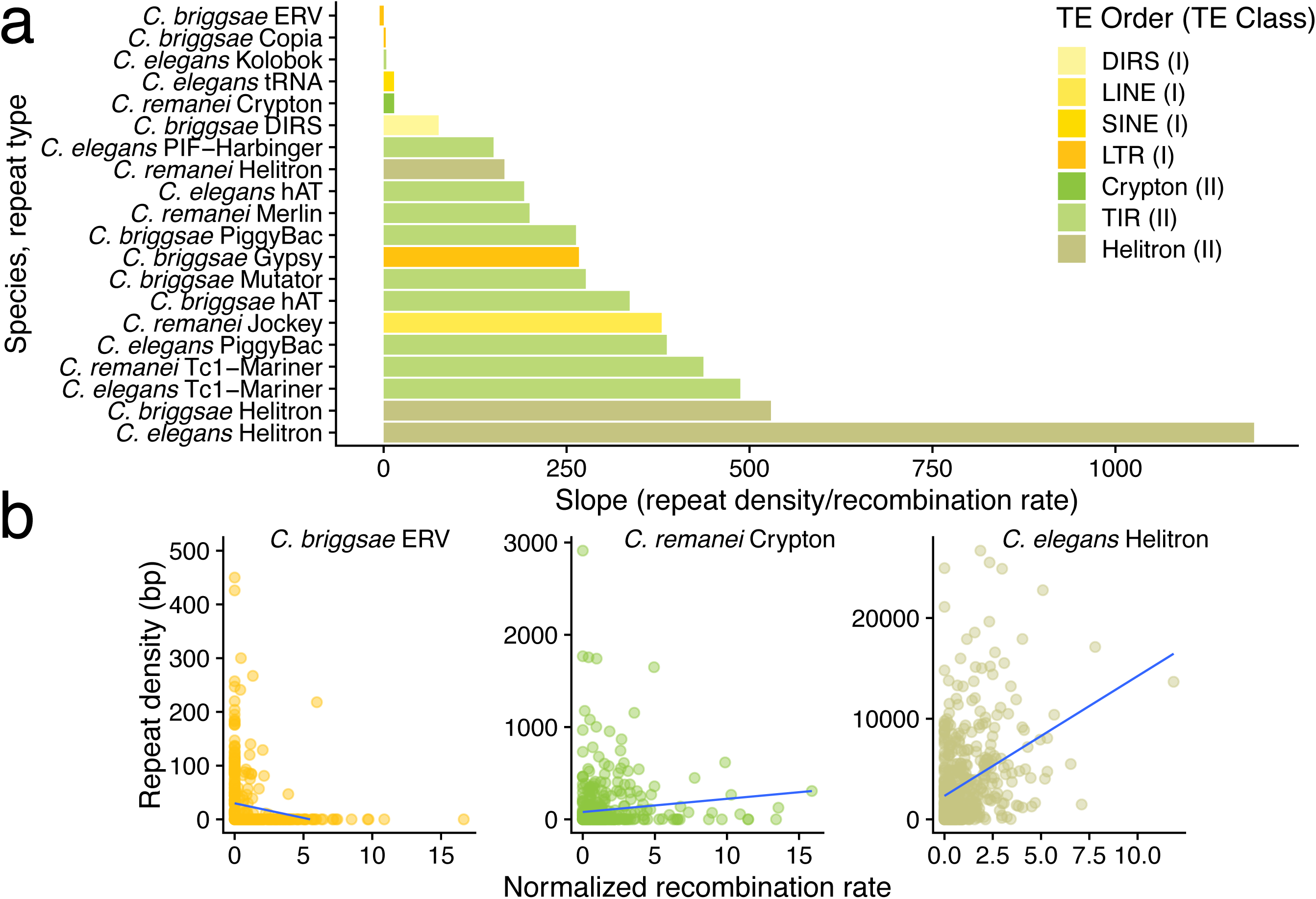
Recombination rate/repeat density relationships vary by repetitive element type and species. (a) All repeat taxa with significant associations with recombination rate. Positive slopes represent positive relationships between repeat density and recombination rate. (b) Three representative scatterplots displaying significant repeat/recombination associations revealed in the previous panel.

## Results

### Conservation of recombination rate structure is chromosome-dependent in C. inopinata

Many multicellular organisms reveal a negative association between recombination rate and repeat density (Fig. 1a). *Caenorhabditis* nematodes represent an atypical exception to this pattern, wherein repeat density is generally positively correlated with recombination rates (Fig. 1a-b). In contrast to most members of the genus, *C. inopinata* harbors uniform repetitive genomic landscapes (Fig. 1b; (Woodruff and Teterina 2020)). Previous work involving evolutionary simulations suggested that the evolution of recombination rates could explain this divergence of the repetitive genomic landscape in *C. inopinata* (Woodruff and Teterina 2020). However, little is known about chromosome-wide recombination patterns in *C. inopinata* (Woodruff et al. 2024). To address this, we constructed a genetic map of *C. inopinata* using 180 F_2_ recombinant progeny (Fig. 2). *C. elegans* (Cutter et al. 2009; Rockman and Kruglyak 2009) and other *Caenorhabditis* species (Ross et al. 2011; Teterina et al. 2020; Noble et al. 2021; Teterina et al. 2023) harbor characteristically structured chromosomes with low-recombining centers and high-recombining arms (Figures 2a, 2c, 3; Supplemental Table 1 sheet 2). And, indeed, this pattern remains largely conserved in *C. inopinata* (Figures 2a, 2c, 3; Supplemental Table 1 sheet 2). However, not all *C. inopinata* chromosomes reveal evidence of arm-center structured recombination domains. Chromosomes IV (Cohen’s *d* arm-center effect size = 0.013; Wilcoxon rank-sum test W = 1531, adjusted *p* = 0.47) and V (Cohen’s *d* arm-center effect size = 0.051; adjusted *p* = 1) appear to have largely uniform rates of recombination across their spans (Figures 2a, 2c, 3; Supplemental Table 1 sheet 2). This is in contrast to most other *Caenorhabditis* chromosomes for which genetic maps have been published, nearly all of which harbor arm-center recombination domains (Figure 3; Supplemental Table 1 sheet 2). The exceptions to this general pattern include *C. tropicalis*, whose chromosome III harbors a largely uniform recombination landscape (Figure 3; Cohen’s *d* arm-center effect size = 0.013; Wilcoxon rank-sum test W = 1531, adjusted *p* = 0.47). In addition, these divergent recombination patterns in *C. inopinata* do not appear to be associated with drastic forms of segregation distortion, which has been associated with various selfish genetic elements in other *Caenorhabditis* species (Seidel et al. 2008; Ben-David et al. 2017; Ben-David et al. 2021; Noble et al. 2021; Xie et al. 2026). Despite this, most *C. inopinata* chromosomes reveal the conservation of this characteristic chromosomal structuring of recombination domains, consistent with patterns observed among its close relatives (Fig. 3).

### Repetitive elements have been decoupled from recombination rates in C. inopinata

In addition to patterns of recombination, various other genomic features discontinuously vary across chromosomes, including genes, polymorphisms, and repetitive elements. Generally, in *Caenorhabditis*, genes are enriched in low-recombining chromosome centers, whereas repeats and polymorphisms are enriched in high-recombining chromosome arms. While genomic features across *C. inopinata* chromosomes were previously described (Woodruff and Teterina 2020), the genetic map described above enables direct comparisons of recombination rates with genomic features. Moreover, existing datasets generated through previous studies (Rockman and Kruglyak 2009; Thomas et al. 2015; Teterina et al. 2020; Noble et al. 2021; Teterina et al. 2023; Crombie et al. 2024; Woodruff et al. 2024) allow the relationships between recombination and genomic features to be compared among species.

Across autosomal genetic maps and physical positions normalized by chromosome lengths (and partitioned into one hundred equal-sized bins per chromosome), the expected correlations emerge among most *Caenorhabditis* species (Fig. 4; Supplemental Table 1 sheet 3). In four *Caenorhabditis* species, repeat content (Supplemental Table 1 sheet 3; OLS β_1_ = 694-5460, *r^2^* = 0.011-0.20, *F* = 5.5-124, *p* = 6.5×10^-26^-0.019) and genetic diversity (OLS β_1_ =3.0×10^-4^ - 0.0012, *r^2^*= 0.017-0.21, *F* =6.2-134, *p* = 1.2×10^-27^-0.013) is positively correlated with recombination rate (Fig. 4). *C. inopinata*, conversely, reveals no significant correlation with recombination rate and repeat content (OLS β_1_ = - 651, *p* = 0.16) or nucleotide diversity (OLS β_1_ =-4.25×10^-7^, *p* = 0.99; Fig. 4). Regarding genes, *C. elegans* (OLS β_1_ =-1.2, *r^2^* = 0.0090, *F* = 4.5, *p* = 0.034; Fig. 4) and *C. briggsae* (OLS β_1_ =-1.5, *r^2^* = 0.014, *F* = 7.1, *p* = 0.0078; Fig. 4) reveal a significant negative relationship between genes and recombination rate. However, *C. remanei* (OLS β_1_ =-0.74, *p* = 0.069), and *C. tropicalis* (OLS β_1_ = - 0.65, *p* = 0.15) reveal non-significant negative relationships with larger slopes and lower *p*-values than those observed in *C. inopinata* (OLS β_1_ =-0.17, *p* = 0.49). Thus, *C. inopinata* is exceptional in that none of these genomic features reveal significant linear relationships with recombination rate in this analysis. A similar pattern emerges when such relationships are examined across individual chromosomes (Fig. 5; Supplemental Table 1 sheet 4). No *C. inopinata* chromosome harbors a significant feature/recombination correlation (Fig. 5, Supplemental Table 1 sheet 4). Conversely, all other *Caenorhabditis* species reveal at least one chromosome with a significant correlation in the expected direction for the given genomic features (Fig. 5, Supplemental Table 1 sheet 4). Here, the only exception was *C. tropicalis*, which reveals no correlation between gene density and recombination rate for any chromosome (Fig. 5, Supplemental Table 1 sheet 4). Above, we showed *C. tropicalis* chromosome III reveals no arm-center difference in recombination rate (Fig. 3). Moreover, *C. tropicalis* in general reveals lower arm-center effect sizes for recombination rates compared to *C. elegans*, *C. briggsae*, and *C. remanei* (Supplemental Table 1 sheet 2). This may explain the absence of feature/recombination relationships seen in *C. tropicalis*, as its recombination rate differences among chromosome domains are weaker. Regardless, taking into account all chromosomes and species, recombination arm-center effect sizes are correlated with diversity, genic, and repeat arm-center effect sizes (Figure 6-7). Taken together, this reveals that the previously-described divergent genomic features of *C. inopinata* (Woodruff and Teterina 2020; Woodruff et al. 2024) are situated within a partially conserved recombination landscape (Fig. 2-3), resulting in the decoupling of recombination rates and genomic features in this species.

### Repeat/recombination associations vary across repeat types and species

Transposable elements are incredibly diverse, harboring an array of replication strategies and sequence features that enables their taxonomic classification (Wicker et al. 2007). Previous studies across systems have subsequently interrogated whether the relationship between repetitive elements and recombination depends on the type of element under consideration (Duret et al. 2000; Kent et al. 2017). We previously examined the chromosomal distribution of repetitive elements across the repeat taxonomy in nematodes; for instance, we found that the uniform repetitive genomic landscape of *C. inopinata* was driven by four taxonomically-unrelated repeat superfamiles (Woodruff and Teterina 2020).

To assess the possibility that repeat-recombination correlations depend on repeat type, we joined our estimates of recombination rates to the repeat taxonomy developed in our previous study ((Woodruff and Teterina 2020); *C. tropicalis* was not included in this work). Out of 26 repeat superfamilies, 15 were found to be significantly correlated with recombination rate in a given species (Fig. 8; Supplemental Table 1 sheet 5). These spanned repeat types harboring a diversity of replication strategies including Type I non-retrotransposons (e.g., SINE tRNAs in *C. elegans;* Fig. 8a), Type I retrotransposons (e.g., Gypsy in *C. briggsae*), and Type II transposons (Tc1/Mariner in *C. elegans* and *C. remanei*). Only one repeat superfamily in one species revealed a negative relationship with recombination rate (ERV in *C. briggsae*; Fig. 6a-b; OLS β_1_ =-5.4, *r^2^* = 0.032, *F* = 17, adjusted *p* = 3.3 x 10^-4^). All other significant repeat-recombination rate correlations were positive (Fig. 8a-b). In addition, coefficients of determination were generally low across these repeat types in the three species (*r^2^* = 0.01-0.11). Taken together, although associations between recombination rate and specific repeat taxa can be identified, it is difficult to predict the density and distribution of repetitive elements based on repeat features and replication strategies.

## Discussion

*The chromosomal distribution of recombination domains is partially conserved in* C. inopinata Here, we generated a genetic map (Fig. 2) to understand *C. inopinata*’s exceptional repetitive genomic landscape (Fig. 1b). Previously, we performed evolutionary simulations that showed that uniform recombination landscapes promote uniform transposable element landscapes (Woodruff and Teterina 2020).

Thus, the intention of this work was to test the hypothesis that *C. inopinata* harbors a uniform recombination landscape, which could explain its unusual chromosome distributions of repetitive elements. Our work reveals both conserved and divergent features of the *C. inopinata* recombination landscape (Fig. 2, 6-7). Three autosomes and the X chromosome reveal higher recombination rates on chromosome arms (Fig. 2, Supplemental Table 1 Sheet 2). Thus, in this respect, most of the *C. inopinata* genome reveals a broadly conserved recombination landscape in comparison to its close relatives (Fig. 2; (Rockman and Kruglyak 2009; Noble et al. 2021; Teterina et al. 2023; Parée et al. 2024)). However, chromosomes IV and V reveal no evidence of recombination domains across chromosome centers and arms (Fig. 2; Supplemental Table 1 Sheet 2). In addition, the magnitude of arm-center differences in recombination are much smaller in *C. inopinata* in comparison to most members of the genus (Fig. 6-7; Supplemental Table 1 Sheet 2). Thus, while qualitative recombination landscape trends persist for most of the genome, there are quantitative and chromosome-specific reductions in recombination rate structure that may in part explain the distribution of genomic features in *C. inopinata*. Specifically, both genetic diversity *and* gene density likewise reveal lower arm-center effect sizes in *C. inopinata* compared to its close relatives (Fig. 6-7; Supplemental Table 1 Sheets 6 & 8). Indeed, gene (Woodruff and Teterina 2020) and polymorphism (Woodruff et al. 2024) chromosomal patterns were previously described. The genic pattern was previously interpreted as a *potential* driver of uniform repetitive genomic landscapes, as uniform selection against TEs was shown to generate such uniform TE landscapes in forward evolutionary simulations (Woodruff and Teterina 2020). Conversely, the reduced arm-center difference in genetic diversity in *C. inopinata* was interpreted to reflect its reproductive mode (Woodruff et al. 2024). That is, as there is overall more effective population-level recombination in the gonochoristic *C. inopinata* (and LD is markedly lower in this species), the impact of recombination domains on diversity was interpreted to be dampened (Woodruff et al. 2024). Our results here offer evidence for an alternative, additional explanation for these patterns. Simply, a reduction in the extent of recombination rate differences among chromosome arms and centers has promoted a reduction in the intrachromosomal differentiation in other genomic features (such as genes, polymorphisms, and repetitive elements; Fig. 6-7; Supplemental Table 1 Sheets 6-8).

What could drive such changes in recombination rates? The evolution of recombination rates has received a considerable amount of theoretical attention (Dapper and Payseur 2017). Following this previous review (Dapper and Payseur 2017), numerous theoretical drivers of recombination rate evolution have been posed. For instance, negative epistatic interactions among beneficial alleles have been proposed to increase recombination rates (Otto and Feldman 1997; Barton and Charlesworth 1998; Kouyos et al. 2007). Presumably, if such alleles emerged along the *C. inopinata* lineage in chromosome centers, this may promote the reduction in recombination rate heterogeneity in this species. Additionally, temporal and spatial heterogeneity in selection regimes have likewise been proposed to be alleviated by the evolution of increased recombination rates (i.e., the reduction in LD among alleles with different effects across space and time enable flexible evolutionary responses to such changing conditions; (Charlesworth 1976; Barton 1995; Otto and Michalakis 1998; Lenormand and Otto 2000; Ortiz-Barrientos et al. 2016; Reeve et al. 2016)). The extent of such environmental heterogeneity in *C. inopinata* is unclear-- while many speculative scenarios could be imagined, why and how this would lead to such specific differences in the fig-and fig wasp-associated *C. inopinata* and *not* its other, more cosmopolitan close relatives remains wholly elusive. The evolution of increased recombination rates has been posed to occur in the face of genetic drift to enable the disassociation of linked deleterious alleles (Hill and Robertson 1968; Otto and Barton 2001; Barton and Otto 2005; Keightley and Otto 2006). This may be consistent with the biology of *C. inopinata*; specialists are expected to have lower effective population sizes and low genetic diversity (the specialist-generalist variation hypothesis (Li et al. 2014)). Thus, a history of bottlenecks or population size reductions may have led to a correlated change in recombination that has persisted and has led to the flattening of the *C. inopinata* genome (if there are alleles capable of modifying recombination rates). Future population genomic and forward evolutionary simulation studies will be needed to properly interrogate and disentangle these possibilities.

The potential proximate causes of such recombination landscape change likewise remain unclear. There is at least one gene, *rec-1*, that has been shown to homogenize recombination landscapes when perturbed in *C. elegans* (Zetka and Rose 1995). As *rec-1* appears to be a *C. elegans*-specific gene, there are not strong, comparable hypothetical molecular drivers of recombination rate change in other *Caenorhabditis* species (like *C. inopinata*). Notably, a number of low-copy chromosomal segregation factors may have been highly duplicated specifically in *C. inopinata* (Supplemental Figure). *icp-1* is an INner CEtromere Protein (Incep) homolog that is critical for chromosome segregation and cytokinesis (Kaitna et al. 2000), and ICP-1 localizes to meiotic spindles (Kaitna et al. 2002; Romano et al. 2003). *C. inopinata* potentially harbors at least seventeen copies of this gene whereas *C. elegans* has one (Supplemental Figure 1) or two (Kaitna et al. 2000). Additionally, *baf-1* encodes a DNA-binding protein that is likewise implicated in chromosome segregation-- *baf-1* loss-of-function mutants harbor chromosome segregation defects (Zheng et al. 2000; Gorjánácz et al. 2007). *C. inopinata* potentially harbors at least eleven copies of this gene whereas *C. elegans* has at least one (Supplemental Figure 1). Future molecular genetic studies aimed at comparing specific factors potentially driving variation in meiotic and crossover phenotypes will be needed to further explore such hypotheses.

*Transposable elements have become decoupled from recombination in* C. inopinata

Despite the apparent loss of recombination domains in two autosomes (and the overall reduction in recombination heterogeneity across chromosomes; Fig. 2, 6-7), the distribution of repetitive and transposable elements is far more uniform in *C. inopinata* than other genomic features (Fig. 6-7), and these patterns span all chromosomes (Fig. 6-7). Thus, while changes in the extent of recombination within chromosomes (and the loss of domains on some chromosomes) may explain some reduction in repeat landscape heterogeneity, it cannot entirely explain these repetitive element distributions. What else could be driving these patterns?

If TEs are largely deleterious by disrupting gene function following insertion, then negative selection against TEs may be a major driver of repetitive genomic landscapes (Charlesworth and Langley 1989; Kent et al. 2017). That is, the genomic landscape of repeats could be caused by the existing genomic landscape of protein-coding genes. As selection would remove TEs more frequently in gene-dense regions, this would drive a negative correlation between gene and repeat density. As there are more genes in chromosome centers (Fig. 6-7), arm-biased repeat landscapes are shaped accordingly. Indeed, evolutionary simulations confirm such intuitions — TEs do not accumulate in genomic regions associated with higher selection coefficients (Woodruff and Teterina 2020). Thus, as *C. inopinata* has fewer genes enriched in chromosome centers than other *Caenorhabditis* species (Fig. 6-7), these gene distributions may have contributed to the uniform repetitive genomic landscape of *C. inopinata*. Together with reductions in recombination rate heterogeneity, this may be sufficient to generate such repeat landscapes. However, there is some indication that some of our data counters this model, as there are *C. tropicalis* chromosomes with little coding gene heterogeneity despite the persistence of repetitive genomic landscapes (Chromosomes I, II, and X; Figure 7; Supplemental Table 1 Sheet 8). Regardless, this scenario suggests that changes in the chromosomal distribution of genes precedes the degradation of repetitive genomic landscapes. Alternatively, existing patterns of chromatin organization (and/or nuclear 3D organization), may likewise play a role in driving recombination landscape evolution. Regardless, a broader phylogenetic scope of nematodes, including the analysis of a large number of chromosome-contiguous assemblies in a phylogenetic comparative methods context, would be required to better test the hypothesis that changes in gene landscapes precede changes in repetitive genomic landscapes.

Although in selection regimes across chromosomes could alter repetitive genomic landscapes, radical alterations of the molecular machinery that regulate transposable elements could also contribute to downstream changes in genomic organization. Small RNAs regulate transposons in multiple systems (Tóth et al. 2016). In *C. elegans*, the argonaut ERGO-1 and its affiliated siRNAs have been shown to regulate transposable elements in oocytes (Fischer et al. 2011; Fischer and Ruvkun 2020). *eri-6/7* encodes a helicase that likewise promotes the formation of siRNAs targeting LTR retrotransposons (Fischer and Ruvkun 2020). And, *eri-9* encodes a DICER-interacting protein that is required for endogenous siRNA expression (Pavelec et al. 2009). *C. inopinata* has lost *ergo-1*, *eri-6/7*, and *eri-9* (Kanzaki et al. 2018). Thus, it is then possible that drastic changes in small RNA regulatory machinery contribute to the evolution of atypical repetitive genomic landscapes. Moreover, as connections between small RNA regulation, epigenetic modifications, and recombination have been demonstrated in other systems (Lee 2015; Lee and Karpen 2017; Huang et al. 2022; Huang et al. 2025), it also remains possible that the loss of endogenous siRNA regulators may contribute to the evolution of recombination rates as well as repetitive genomic landscapes (Huang et al. 2025). The multiple, concurrent changes in gene density, recombination, TE density, and diversity make disentangling the causes of genomic organization difficult. Despite this, situating existing mutations (such as those in the recombination machinery (such as *rec-1*, (Zetka and Rose 1995; Parée et al. 2024)) and siRNA machinery (such as *ergo-1*, (Fischer and Ruvkun 2020))) into experimental evolution contexts may afford ways to test hypotheses regarding the causes and consequences of the evolution of repetitive elements and recombination rates (Teotónio et al. 2017).

### No clear connection between repeat type distribution and recombination rate distribution

As different kinds of TEs have different modes of replication (Wicker et al. 2007; Wells and Feschotte 2020), different TE types may be expected to harbor variable responses to evolutionary forces such as recombination. Indeed, previous findings have found that the relationship between recombination and repeat density varies by repeat type (Duret et al. 2000; Wright et al. 2003; Tian et al. 2009; Daron et al. 2014). Here, we find only fifteen repeat superfamilies (out of 26 found in our genomes) to harbor any significant relationship with recombination rate (Fig. 8). Many of these superfamilies reveal such a relationship in only one (11 superfamilies) or 2-3 (4 superfamilies) nematode species (Fig. 8). The Helitron repeat superfamily was most frequently correlated with recombination (associated with recombination rates in three species; Fig. 8; Supplemental Table 1 Sheet 5). These superfamilies spanned a wide range of TE structural and replicative diversity (Fig 8.; Supplemental Table 1 Sheet 5). Despite this, Type II DNA transposon superfamilies tended to have stronger positive relationships with recombination rates (Fig. 8). However, Type I retrotransposons were also shown to be correlated with recombination in some cases (Fig. 8). Regardless, a similar pattern wherein a diverse set of repetitive elements were associated with chromosome arms was previously observed (Woodruff and Teterina 2020). Thus, despite evidence of replicative strategy being connected to recombination in other systems (Kent et al. 2017), this pattern is not consistently seen here. At least in *Caenorhabditis* nematodes, the fine structure of recombination does not necessarily or consistently predict the accumulation of any specific type of transposable element.

### An unexpected relationship between recombination and repetitive elements in nematodes

Across a broad range of taxa, TE abundance is generally negatively correlated with recombination rate (Fig. 1; (Kent et al. 2017)). As has long been known (Cutter et al. 2009), nematodes reveal a positive correlation among these traits, cutting against this apparently widely conserved pattern (Fig. 1). The broad patterns observed in most taxa often reveal the accumulation of TEs in low-recombining centromeric regions (Slotkin and Martienssen 2007; Plohl et al. 2014; Termolino et al. 2016). *Caenorhabditis* nematodes harbor holocentric centromeres, with multiple sites of kinetocore attachment and no obviously segregated (or clearly delineated) centromeric regions on its chromosomes (Albertson and Thomson 1982; Albertson and Thomson 1993). If centromeric functions are the major drivers of TE accumulation in systems with monocentric centromeres, this may explain why nematodes are so unusual in their repetitive genomic landscapes. As their chromosomes have no large centromeric landing pages, repetitive elements accumulate instead in otherwise gene-poor regions, regardless of how frequently they recombine. If chromosome structure drives the relationship between recombination and TE abundance, it would be predicted that this expected negative relationship breaks down across taxa with holocentric chromosomes. Holocentricity is not rare — multiple clades of green algae, angiosperms, arachnids, myriapods, and insects have holocentric centromeres (Mandrioli and Manicardi 2020). A systematic, phylogenetically informed investigation into the relationship between recombination rates, repetitive element density, and centromeric structure will be required to test this hypothesis. Regardless, the breakdown of the connection between recombination and TE abundance in *C. inopinata* shown here, in tandem with the long-standing positive correlation between these traits in other *Caenorhabditis* species (THE C. ELEGANS SEQUENCING CONSORTIUM 1998; Stein et al. 2003; Fierst et al. 2015; Woodruff and Teterina 2020; Noble et al. 2021; Stevens et al. 2022; Teterina et al. 2023), demonstrates that there is more at play in driving the evolution of genomic organization than recombination alone. Proximate molecular drivers (such as centromeric organization and small RNA pathways), as well as historically-contingent modulators of selection (such as gene density heterogeneity) likely jointly contribute to the diverse distributions of repetitive elements seen in this system. An exception such as *C. inopinata* offers a unique opportunity for future studies examining these myriad drivers of genome evolution.

## Data archiving

Whole-genome sequencing reads have been submitted to the NCBI SRA (under BioProject ID PRJNA1471261). Other data and code have been deposited in Github (https://github.com/gcwoodruff/C_inopinata_genetic_map_2026/).

## Acknowledgments

We would like to thank Jeff Bishop and the University of Oregon GC3F Genomics Core Facility for preparing Illumina libraries and performing whole-genome sequencing for this study. Some of the computing for this project was performed at the OU Supercomputing Center for Education & Research (OSCER) at the University of Oklahoma. We thank Anastasia Teterina for providing comments on an earlier version of this manuscript.

## Funding

This work was funded in part by an award from the National Science Foundation to GCW (Award #2238788). This work was supported in part by funding from the University of Oklahoma.

## Author contribution statement

Kimberly A. Moser: Investigation, Resources, Writing – review and editing | Courtney King: Investigation, Resources | Gavin C. Woodruff: Conceptualization, Data curation, Formal analysis, Funding acquisition, Investigation, Methodology, Project administration, Resources, Writing – original draft, Writing – review and editing.

## Conflict of interest

The authors declare no competing financial interests.

## Research ethics statement

This work describes observations of *Caenorhabditis* nematodes. The study of such nematodes is not associated with any required ethical considerations connected to the study of humans, vertebrates, or higher invertebrates.

## Supplementary data

Supplemental Figures and Tables can be found at *Journal*.

## Notes

### Competing Interest Statement

The authors have declared no competing interest.

### Summary of Updates

The original post was missing Figure 5. Additionally, some main text language was obscured by figure boxes. These issues have hopefully been addressed.

