## Supplemental Figure for "Recombination and repetitive genomic landscapes are decoupled in a close relative of *Caenorhabditis elegans*"

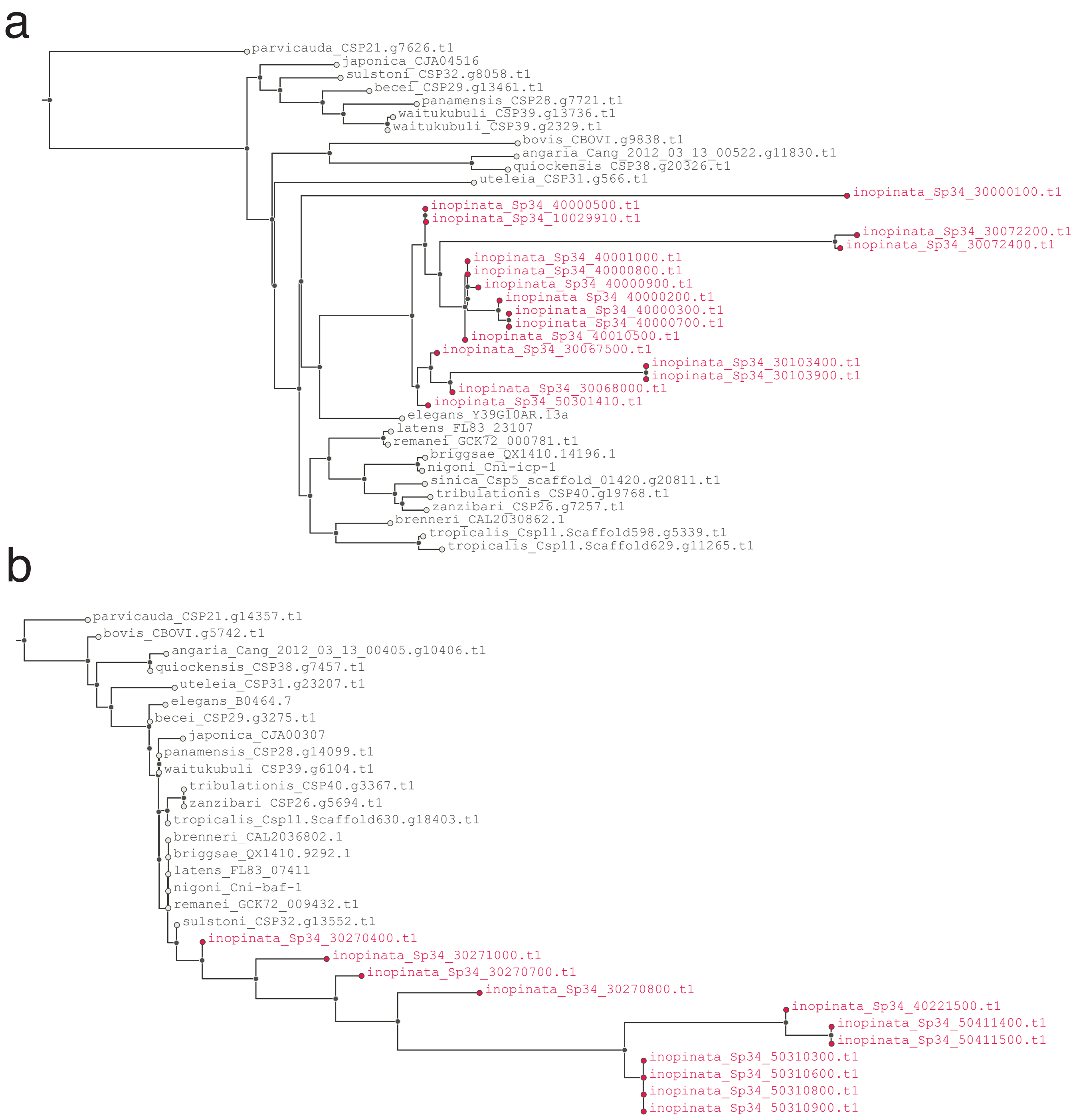


Supplemental Figure 1. *C . inopinata* harbors more copies of specific chromosome segregation machinery genes than other *Caenorhabditis* species. (a) *ce-icp-1* orthogroup; (b) *baf-1* orthogroup. The OrthoFinder software placed one *C. elegans* copy of these genes in these clusters, whereas 17 (*icp-1* orthogroup) and eleven (*baf-1* orthogroup) *C. inopinata* genes (in red) were placed in these groups. Gene trees were generated with FastTree following orthogroup inference with OrthoFinder; bootstrap values were not collected.
